## Supplemental Text and figures for "Inferring control objectives in a virtual balancing task in humans and monkeys"

### 1. Subject-to-subject variability of hand/cursor RMS ratio

In Figure 2, a seemingly deviant behavior was observed for the Hand/Cursor RMS ratio of monkey J compared to the other monkey and the human subjects. This could be due to subject-to-subject variability. Given that data from only two monkeys were available, we examined this possibility by presenting the individual hand/cursor RMS ratio for all individual human subjects. Figure S1 shows that there was indeed variability across subjects, with some not exhibiting a clear trend with task difficulty. However, on average, the RMS ratio showed a slight decrease as trials grew more difficult, as was earlier shown in Figure 2.

### 2. Alternative metrics for inferring control objectives from behavior

We used two main metrics in our analysis of inferring the control objective from behavior based on cursor movement, namely, the mean and RMS of cursor position/velocity (Figure 5 and 6). In Figure 5 we demonstrated that the choice of control objective affected the correlation between cursor mean position and cursor mean velocity in the state space of cursor movement. As one of the reviewers observed, since the cursor mean velocity over a trial determined the cursor final position in that trial, one could interpret the correlation between cursor mean velocity and its mean position in terms of the autocorrelation function (acf) of cursor position.

In particular, under Position Control, the final cursor position would be relatively uncorrelated with the average position, and hence the temporal acf of position would be narrow. In contrast, under Velocity Control, the final position tends to be similar to the average position, and thus the acf would be wider. We explored this insight by calculating the width of the acf of cursor position for 200 simulated trials at four different $\lambda$ values for each control objective. Figure S2A shows example cursor and hand traces, together with the corresponding cursor acf and its width (defined as the width of a rectangle with area equal to the area under the absolute value of the acf). In Figure S2B, the distribution of acf width across trials is compared between Position Control and Velocity Control for four example $\lambda$ values. As expected, given the relation between mean and final position under the two control objectives, the distributions of acf widths separate between the two control objectives. As such, acf width could be another metric to dissociate between different control objectives. However, there is similar overlap between the two objectives, resulting in similarly ‘undecided’ trials as the metrics we used.

Yet another alternative behavioral metric that could potentially differentiate between different control objectives is one that includes both hand and cursor movement, such as the hand/cursor RMS ratio. Figure S3 shows the distribution of hand/cursor RMS ratio across simulated trials, generated based on Position Control or Velocity Control for different $\lambda$ values. As shown, this metric also demonstrates the separation of control objectives, albeit with dependence on $\lambda$: as the task difficulty increased, the distributions begin to converge, thereby becoming less distinguishable (this effect can also be observed in Figure 4A).

Overall, these alternative metrics also reflect the distinction between control objectives in behavior. While they do not offer any observable improvement over our previously used metrics (shown in Figures 5 and 6), it is possible that a more exhaustive examination of behavioral features could lead to metrics that better discriminate between control objectives. Such an investigation is beyond the scope of this study.

### 3. Sensitivity analysis of model parameters

We further investigated whether the distinction into two behavioral patterns could also be accounted for by changing other model parameters, specifically the relative cost of effort, motor noise, or sensory delay. To this end, we conducted a series of simulations wherein the control objective remained fixed at either Position or Velocity Control, but effort cost (***U*** in equation 2), noise magnitude ($\epsilon$ in equation 10) and sensory delay were varied independently.

We first examined whether changing the effort cost under a fixed control objective could account for the variability of behavior across groups in Experiment 2. For each control objective, the effort cost was varied between ***U***=10, ***U***=100, and ***U***=1000, and the resulting change in behavior was examined. As shown in Figure S4A, the overall performance within a given control objective remained independent of effort cost. In particular, effort cost did not affect the distributions of cursor mean (Figure S4B) and cursor RMS (Figure S4C), indicating that the distinctive patterns observed in Experiment 2 could not be explained solely by changing the effort penalty.

Changing the sensory delay time (from 30ms to 70ms; Figure S5) did impact the success rate at a given $\lambda$, not unexpectedly. However, it did not affect the lag, correlation, RMS ratio, or the distributions of cursor mean and cursor RMS. Changing the level of motor noise (from 10% reduction to 10% increase in noise standard deviation; Figure S6) likewise impacted the success rate, but not the other metrics. Overall, these results demonstrated that different control behaviors in the data were predominantly explained by varying the control objective and not effort cost, noise level, or sensory delay.

### 4. Effect of task difficulty on control objectives

We examined whether and to what extent subjects used the same control strategy in different task difficulty levels ($\lambda$ values). We only examined this question in subjects who were instructed to adopt a given strategy, Position or Velocity Control. Figures 5 - 8 presented the data for an ensemble of $\lambda$ values, ranging up to the critical $\lambda$ value ($\lambda_{c}$; associated with 50% success rate). Here, we replot Figure 5 by separating the trials into two clusters based on task difficulty, Easy and Moderate$\lambda$values. The figure shows that the relative behavioral difference between the two control objectives remains qualitatively the same across difficulty levels. Specifically, Figure S7A shows the joint distribution of cursor mean velocity against cursor mean position for Easy ($\lambda$ $\leq$ 70% $\lambda_{c}$), and Moderate (70% $\lambda_{c}$ < $\lambda$ $\leq$ $\lambda_{c}$) conditions. The data are presented for two example subjects, S4 from the Position Control instruction group (brown) and S1 from the Velocity Control group (cyan). As shown, the structure of the data distribution remains approximately the same across Easy and Moderate difficulties, and this is also true with the model simulations (Figure S7B). Importantly, the relative structural difference between the two control objectives, quantified by the correlation coefficient between cursor velocity and cursor position R remains unchanged across different difficulty levels as shown in Figure S7C. In all cases, R was larger for Velocity Control, indicating consistency in control objective across $\lambda$ values.

The effect of time, or practice, on the control objective is subsumed in the analysis above because subjects performed the task progressing from easy to difficult trials: easy trials were performed early in the experiment, and they became increasingly more difficult towards the end of the experiment. To better examine the time course of possible changes in the control objective, we calculated the probability with which a given trial was performed under the Position Control objective. This probability was obtained from the classifier as applied to each trial. Figure S8 shows this probability over the course of trials for each individual in each instruction group. As shown, though the trends were noisy, the probabilities remained generally higher for the Position Control group, and lower for the Velocity Control group as expected. Mainly, subjects’ performance generally remained within the bounds of the control objective they were instructed with throughout the course of experiment.

### 5. Optimal control gains under different control objectives

The optimal feedback controller in our approach calculates the optimal gains that minimize the cost function (Eq. 2) for a given control objective. The resulting control command is $u=-Lx$ (Eq. 12), where $x$ is the state vector and $L$ is the gain vector. Because the cost function depends on the control objective as well as the system dynamics (specifically, the value of $\lambda$), the optimal position and velocity gains will likewise depend on $\lambda$ as well as the control objective. Figure S9 illustrates the control gains for each cursor state for Position Control and Velocity Control across a range of $\lambda$ values. As shown, the optimal gains under the same control objective vary with task difficulty, i.e., $\lambda$. This indicates that as $\lambda$ changes, the gains also need to change in order to (optimally) meet the control objective. Importantly, the choice of control objective is strongly reflected in the cursor position gain, where the two control objectives show opposing trends across task difficulties.

### 6. Perturbation simulations

To explore the potential for perturbation experiments to enhance the ability to discriminate between control objectives, we implemented a random cursor jump (left or right of screen center) at the start of each trial in 1000 simulation trials of Position Control, and of Velocity Control, over a range of difficulty levels. The magnitude of the cursor displacement was randomly sampled from a normal distribution with zero mean (corresponding to screen center) and a standard deviation of 1cm. Figure S10A shows the simulation results for success rate, hand-cursor lag and correlation, and the hand/cursor RMS ratio. As shown, despite the similarity of the success rates for both control objectives, the other metrics show more pronounced differences between the control objectives, compared to the unperturbed simulations (i.e., Figure 4). Interestingly, the difference was more systematic when looking at the cursor states in the mean or RMS spaces. Figure S10B shows the joint distribution of cursor mean position and mean velocity, where the different control objectives showed opposite correlations between cursor position and velocity. Similarly, Figure S10C shows greater separation between the RMS distributions in Position and Velocity Control.

These simulations demonstrate the potential to more robustly differentiate between different control objectives at the behavioral level, and consequently allow for clearer parsing of the neural data to search for neural correlates of control objectives. However, we leave the experimental assessment of these predictions for future studies.

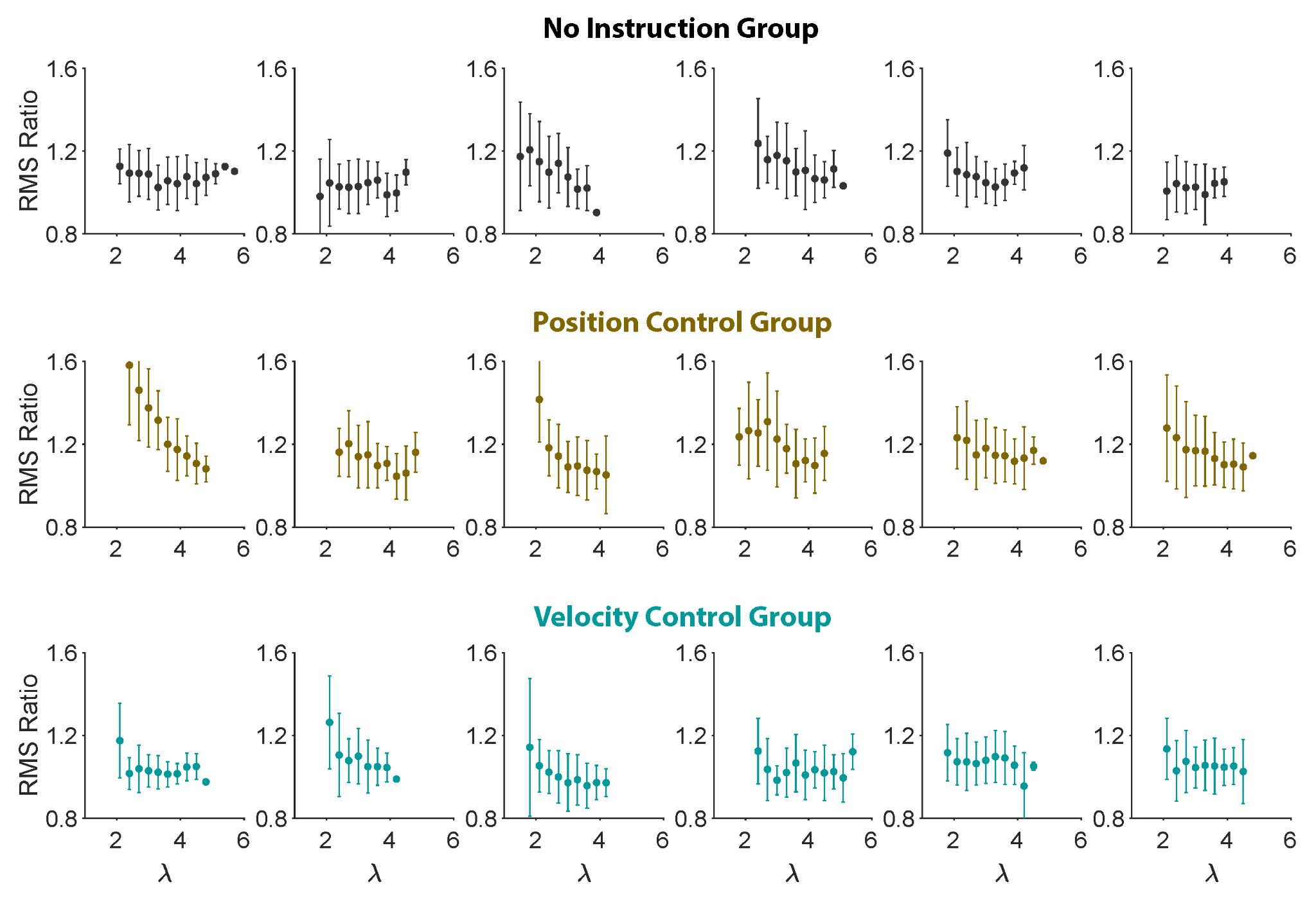

Figure S1: Hand/Cursor RMS ratio for individual participants in all three groups: **Top:** No Instruction group, **Middle:** Position Control group, and **Bottom:** Velocity Control group. The error bars indicate the standard deviations (SD) across trials for each difficulty level.

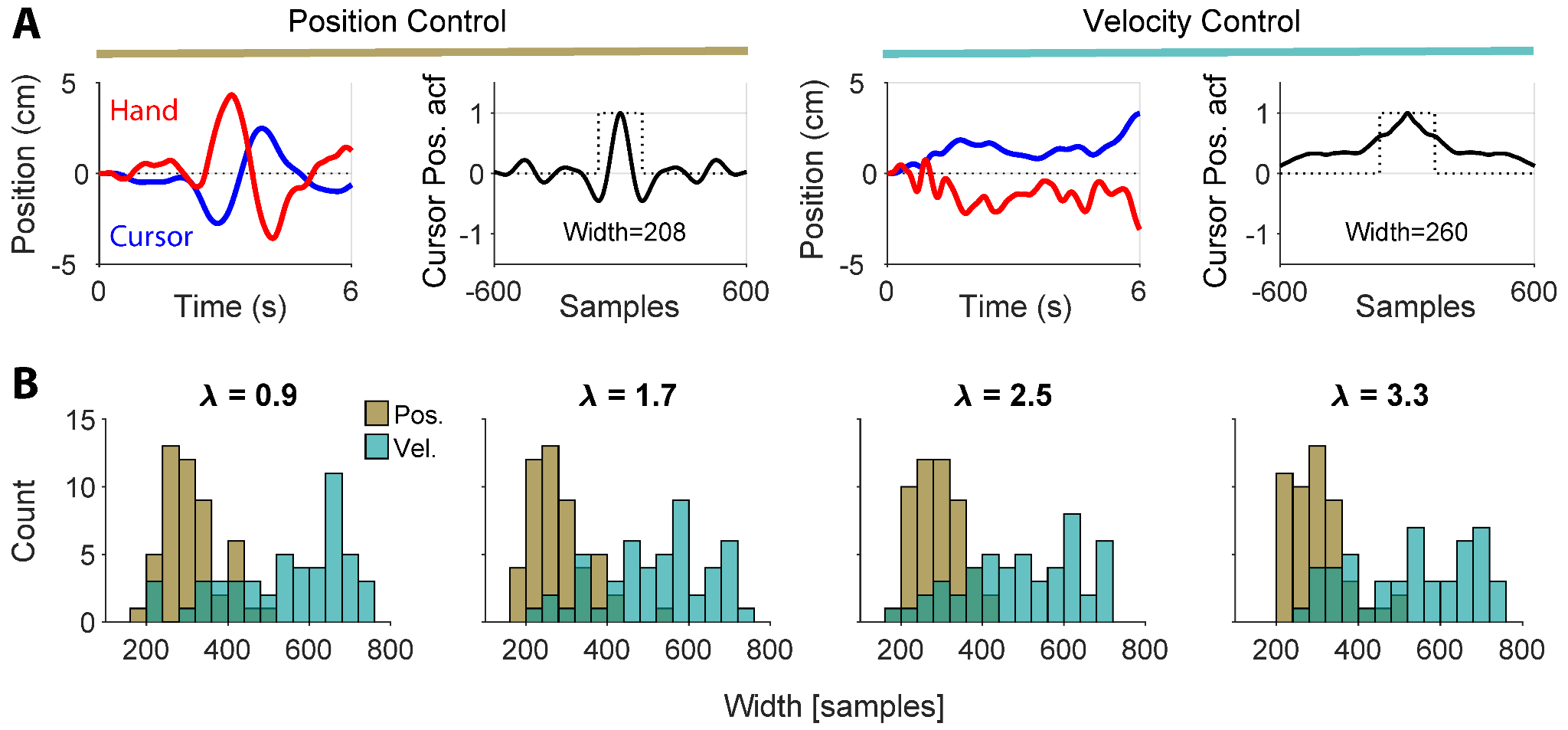

Figure S2: **A**. Sample trials with autocorrelation functions (acf) of cursor position shown for position control (columns 1 and 2), and velocity control objectives (columns 3 and 4). The dotted rectangle in the acf plots shows the acf width (see text for definition). **B**. Histograms of the acf width shown for Position (brown) and Velocity (cyan) control objectives. Each panel shows the results for a different value of $\lambda$ as indicated.

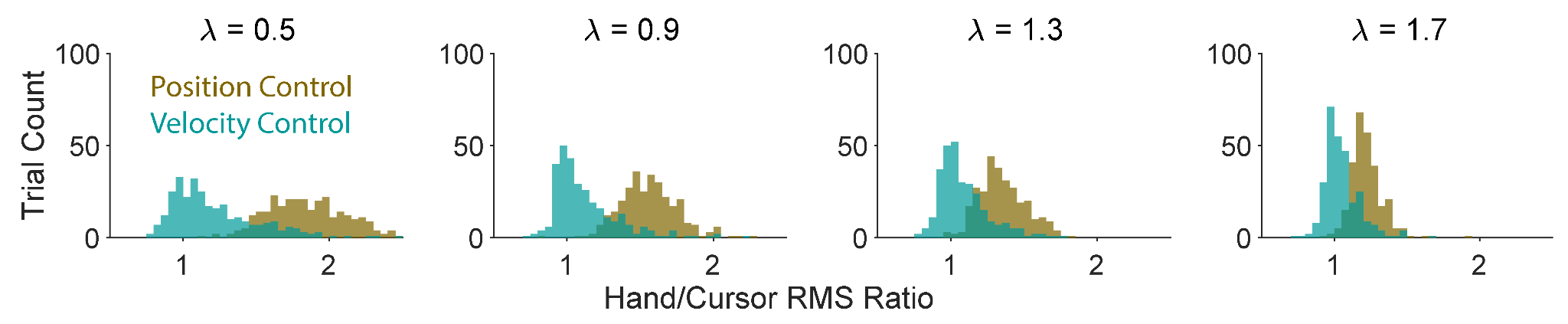

Figure S3: Distribution of hand/cursor RMS ratio over trials, calculated for simulated trials under Position Control (brown) and Velocity Control (cyan), for different $\lambda$ values.

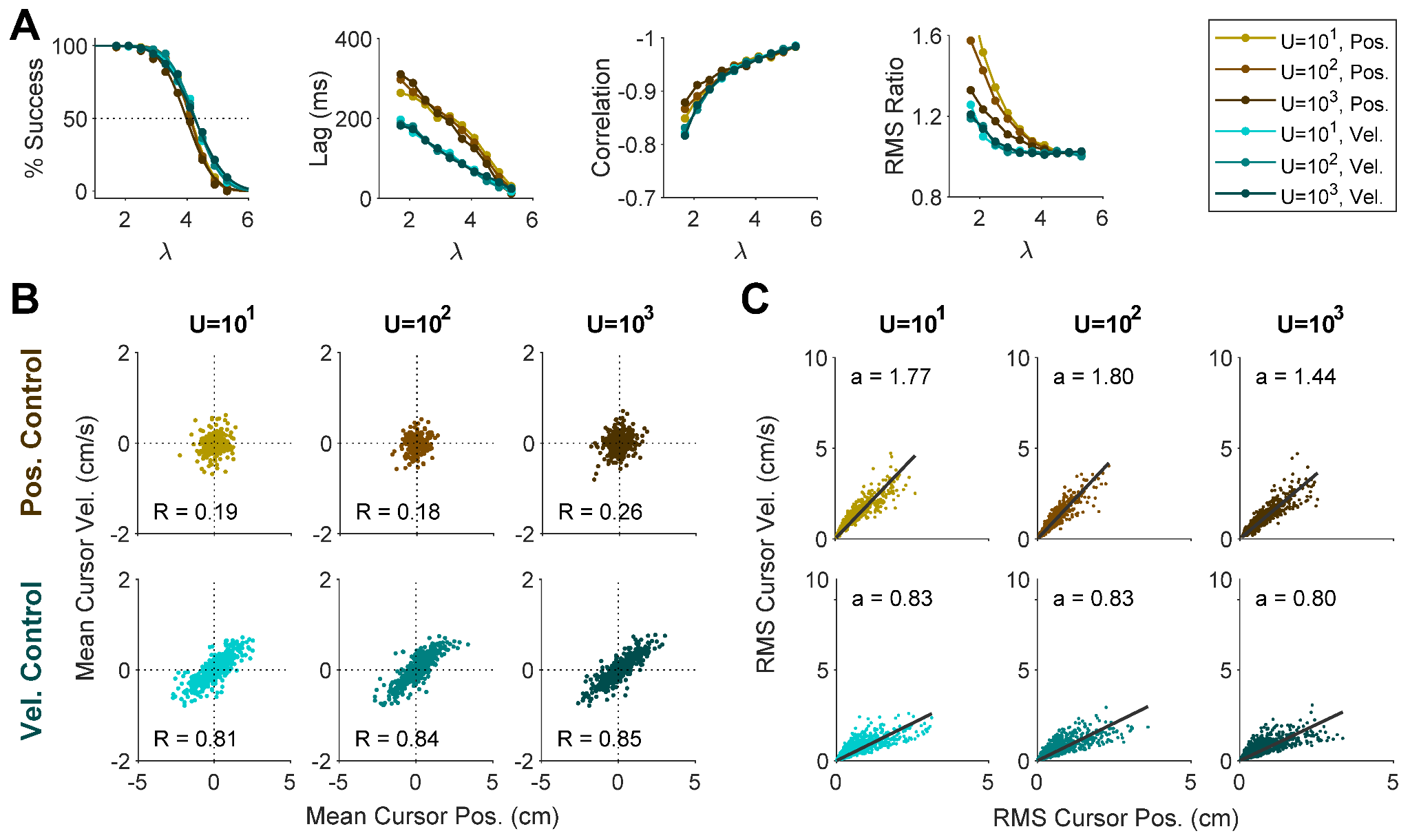

Figure S4: Effect of effort cost on control behavior represented by the: **A**. Aggregate performance measures, **B**. Distribution of mean cursor movement, and **C**. Distribution of RMS of cursor movement.

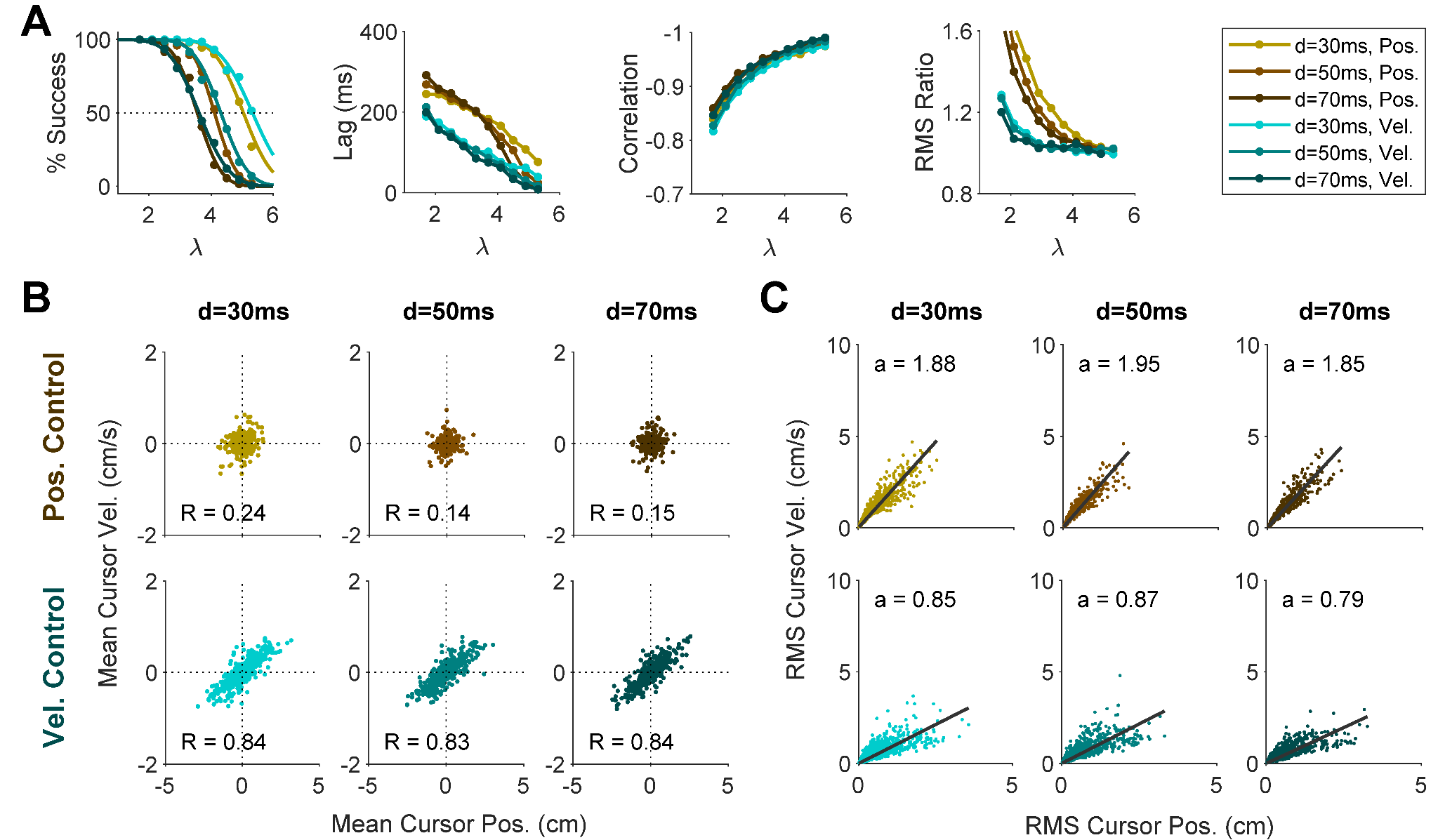

Figure S5: Effect of sensory delay on control behavior represented by the: **A**. Aggregate performance measures, **B**. Distribution of mean cursor movement, and **C**. Distribution of RMS of cursor movement.

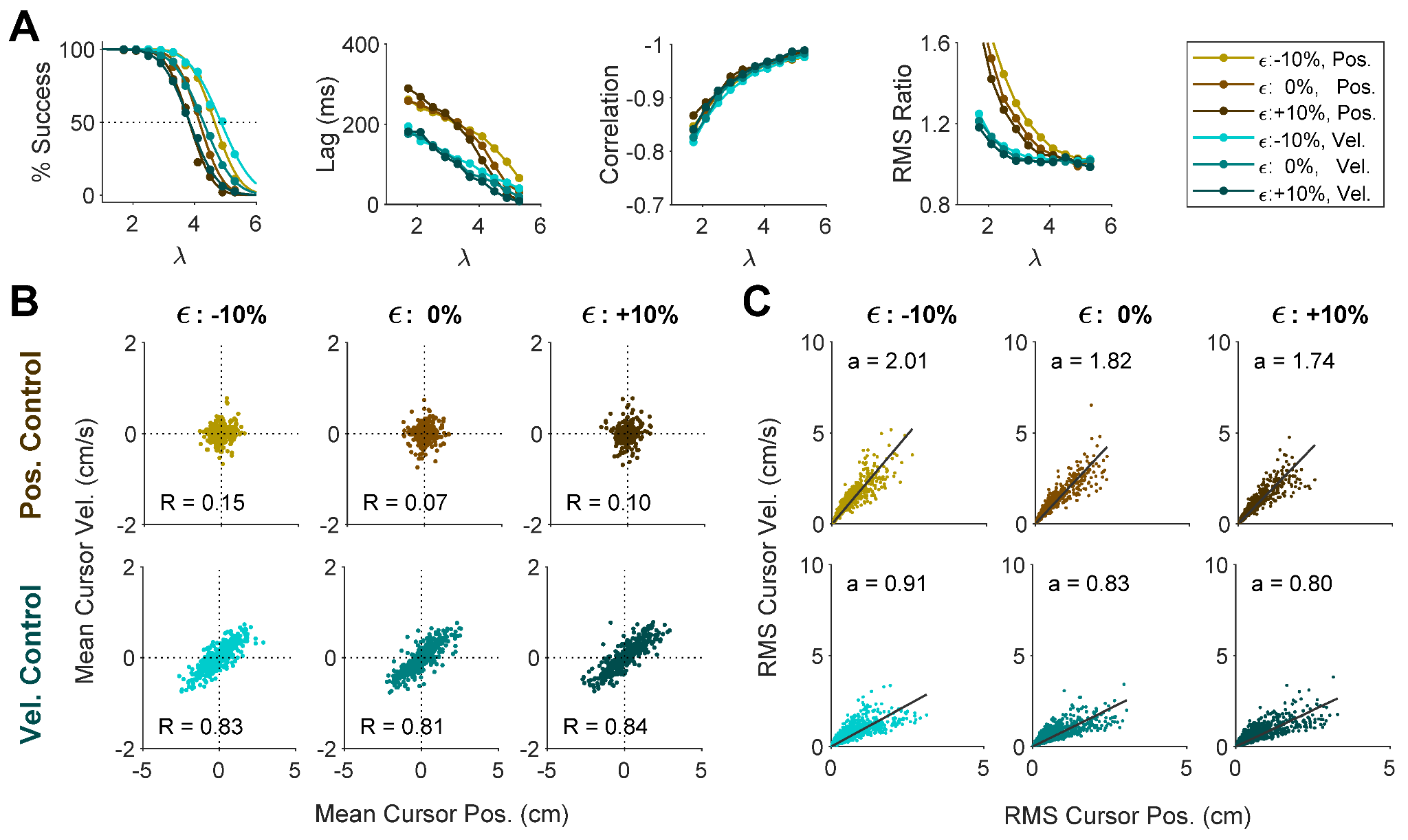

Figure S6: Effect of changing motor noise (10% reduction to 10% increase) on control behavior represented by the: **A**. Aggregate performance measures, **B**. Distribution of mean cursor movement, and **C**. Distribution of RMS of cursor movement.

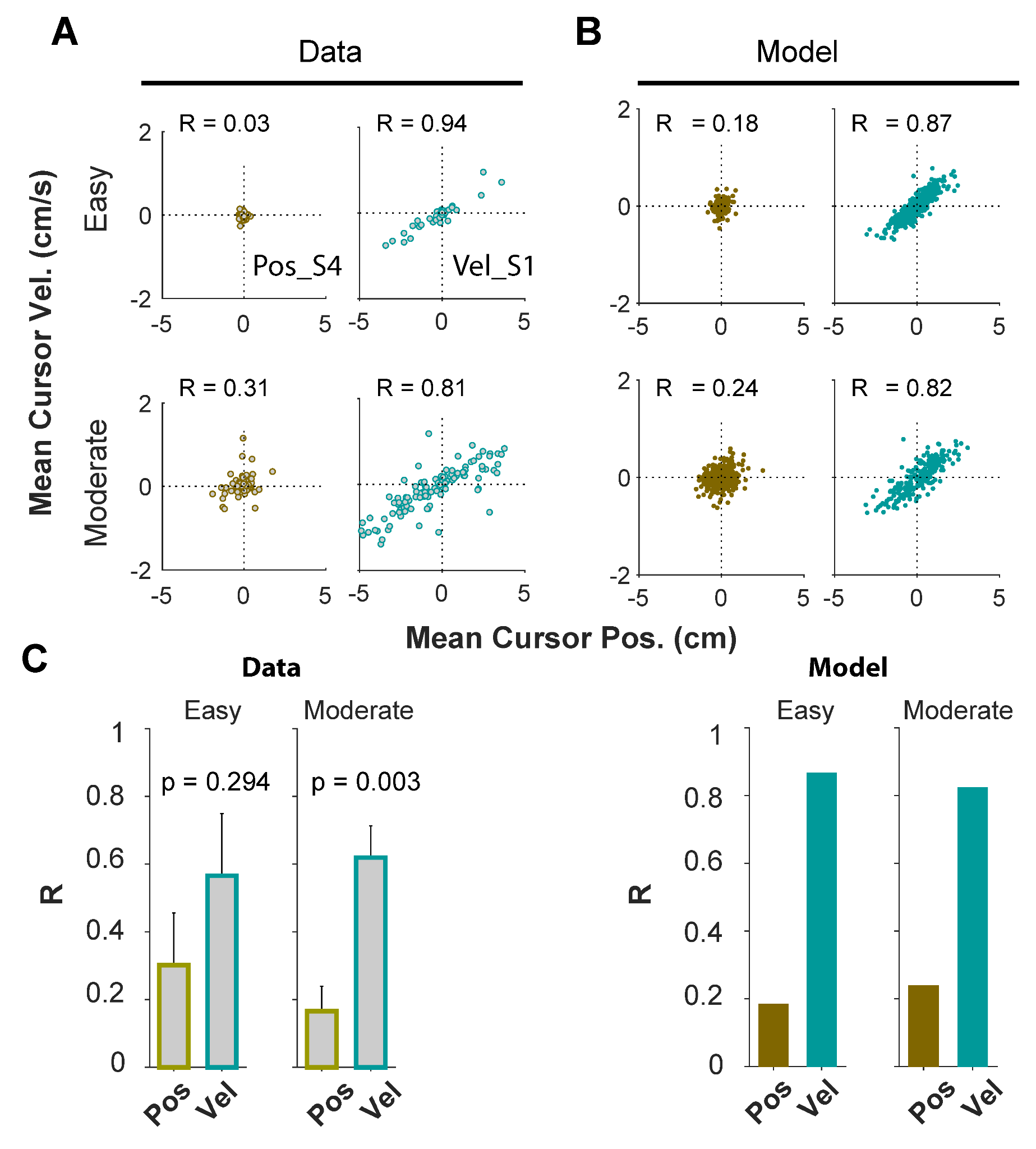

Figure S7: Joint distributions of cursor mean position and cursor mean velocity, shown for experimental data (**A**) and model data (**B**), separated into two difficulty levels: Easy ($\lambda$ $\leq$ 70% $\lambda_{c}$), Moderate (70% $\lambda_{c}$ < $\lambda$ $\leq$ $\lambda_{c}$). **C.** Correlation coefficient (R) between cursor mean position and mean velocity for different difficulty levels, plotted for data (left) and model (right).

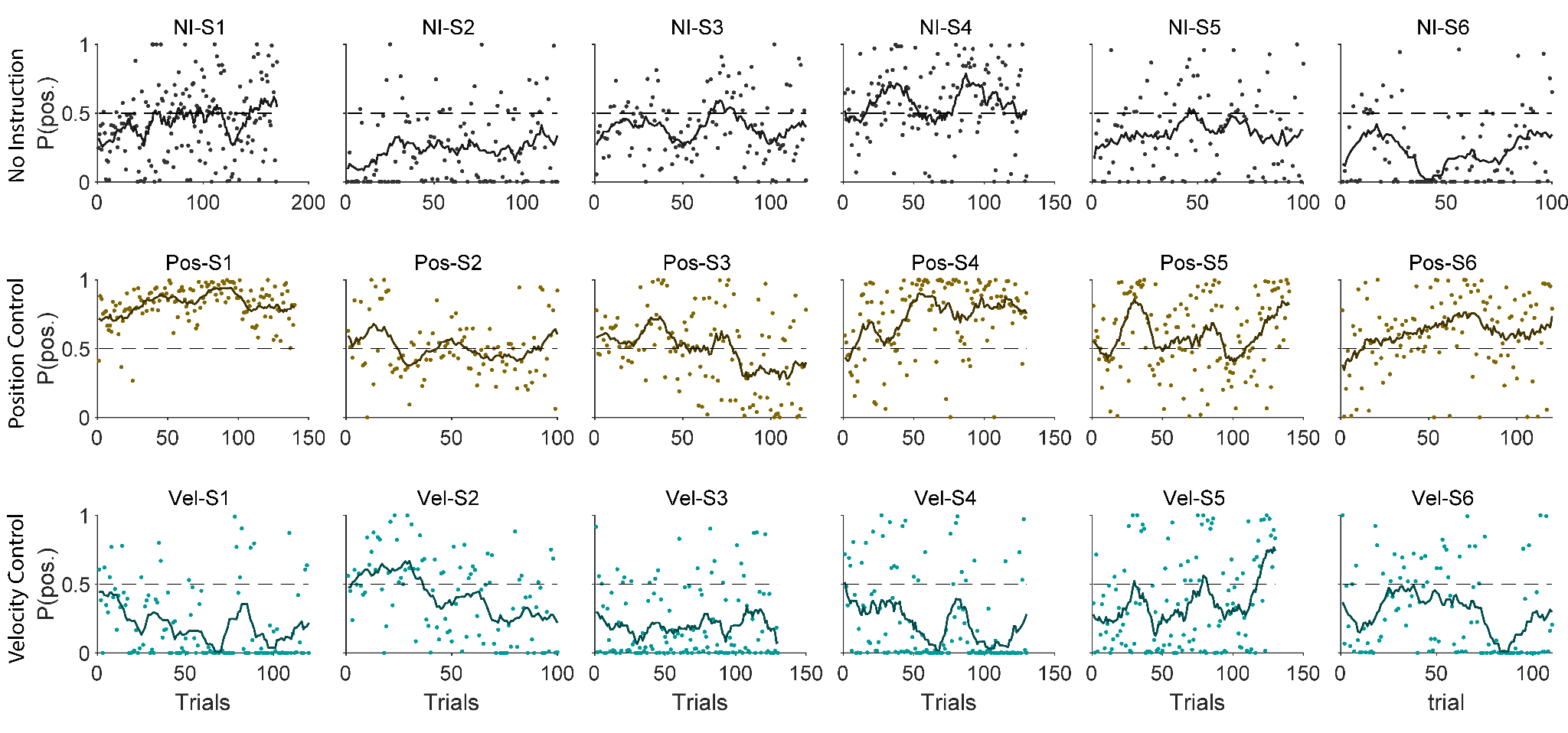

Figure S8: Probability of each trial performed under Position Control for each individual and group (top: Position Control group; bottom: Velocity Control group).

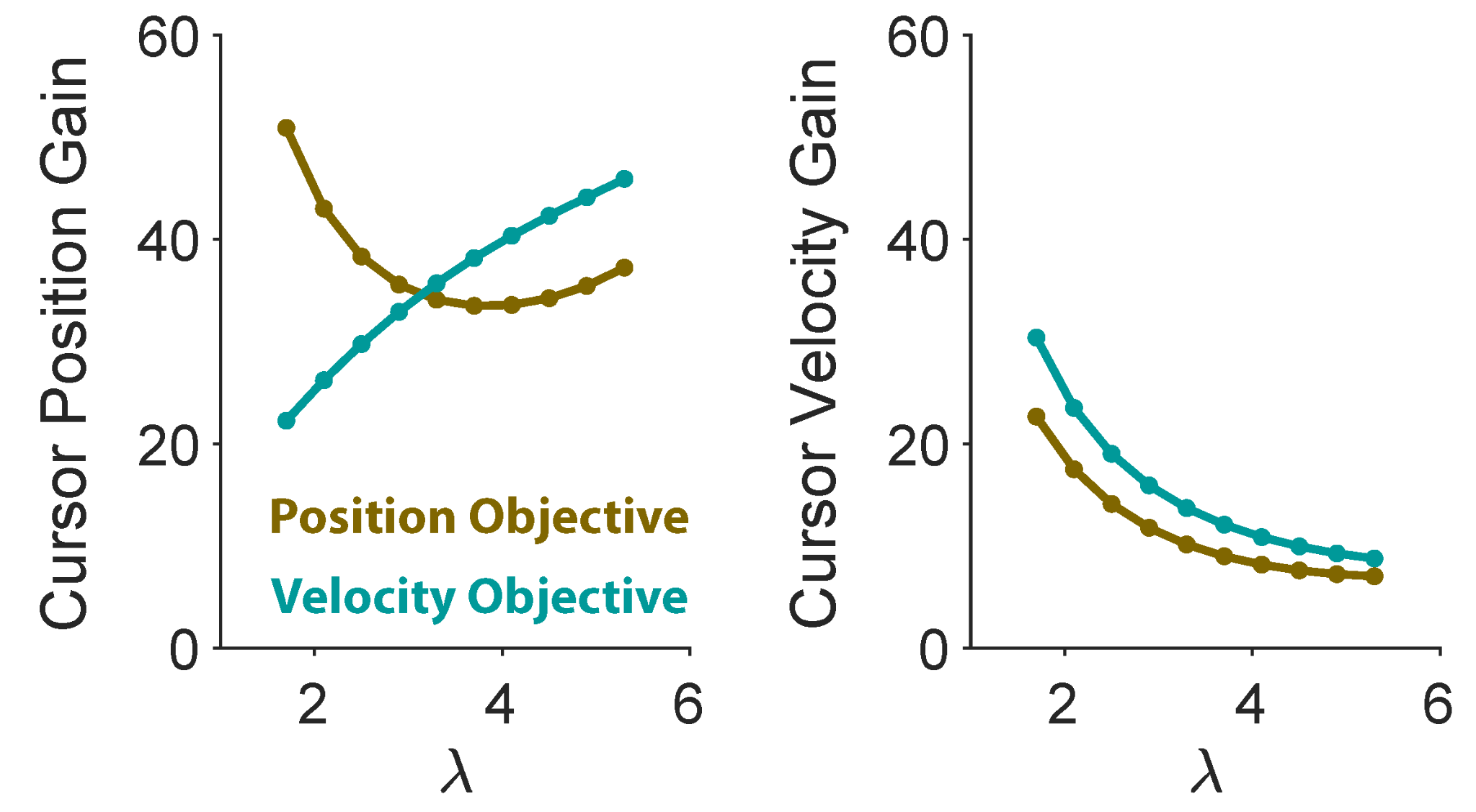

Figure S9: Optimal control gains corresponding to cursor position (left) and cursor velocity (right), obtained under Position Control (brown) and Velocity Control (cyan).

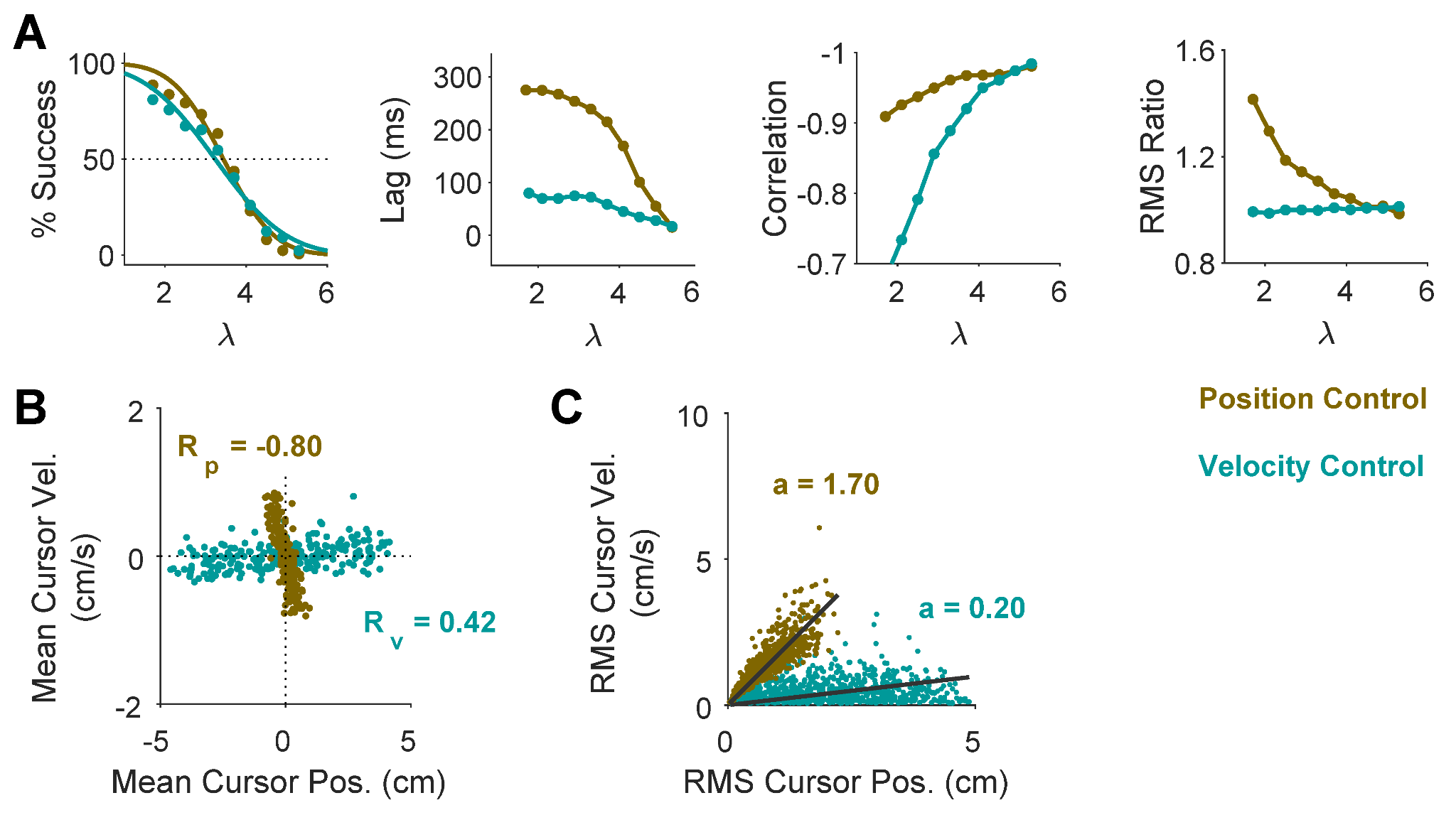

Figure S10. Model simulations of the CST task when introducing perturbations (random cursor jumps) at the start of each trial. **A**. Aggregate performance of success rate, hand/cursor lag, correlation and RMS ratio as a function of difficulty level, shown for each control objective (brown: Position Control; cyan: Velocity Control). **B**. Joint distribution of cursor mean position and cursor mean velocity under different control objectives. **C**. Distribution of cursor RMS position and RMS velocity under different control objectives.
